# An efficient root transformation system for recalcitrant *Vicia sativa*

**DOI:** 10.1101/2020.06.14.151175

**Authors:** Vy Nguyen, Iain R. Searle

**Author notes:** **Correspondence:** Iain R. Searle.

## Abstract

Common vetch (*Vicia sativa*) is a multi-purpose legume widely used in pasture and crop rotation systems. Vetch seeds have desirable nutritional characteristics and are often used to feed ruminant animals. Although transcriptomes are available for vetch, problems with genetic transformation and plant regeneration hinder functional gene studies in this legume species. Therefore, the aim of this study was to develop an efficient and rapid hairy root transformation system for common vetch to facilitate functional gene analysis. We infected the hypocotyls of five-day old *in vitro* or *in vivo,* soil grown, seedlings with *Agrobacterium rhizogenes* and produced transformed hairy roots 28 days later at 24% and 43% efficiency, respectively. Seventy-nine percent of the hairy roots from the *in vitro* plants showed stable expression of a co-transformed marker β-glucuronidase (GUS). In summary, transgenic hairy roots were obtained within 28 days, and are sufficient to facilitate functional gene analysis in common vetch.

## INTRODUCTION

Common vetch (*Vicia sativa*) is a multi-purpose legume crop that is widely used in pastures (Sattell et al., 1998), intercropping and crop rotation regimes. Vetch is grown mainly in Europe, Asia, North America and Oceania, with 54% of the production originating in Europe (FAO data, 1994 - 2017). Vetch seed is rich in protein, up to 32%, and has very low amounts of lipids make it an appealing food source (Mao et al., 2015). Moreover, symbiotic nitrogen fixation of nitrogen from the atmosphere to plant available nitrogen is beneficial in crop rotation systems by reducing the amount of nitrogen fertilizer required (Hargrove, 1986). Vetch also exhibits considerable drought tolerance (Tenopala et al., 2012) which is highly valued as farming in arid areas increases due to climate change (Lobell and Gourdji, 2012).

In recognition that vetch has high potential for genetic improvement, there are several established breeding programs in Europe and Australia. These programs mainly use conventional breeding methods, selecting for traits such as yield improvement (Mikić et al., 2019), non-pod shattering (Abd El-Moneim, 1993), soft seed and rust resistance (GRDC, 2018). Current research focusses on understanding the molecular basis of important agricultural traits, aiming to facilitate molecular plant breeding (Moose and Mumm, 2008). The recent emergence of transcriptome data (Kim et al., 2015;Dong et al., 2017) will accelerate the identification of the genes controlling traits such as pod shattering (Dong et al., 2017), γ-glutamyl-β-cyanoalanine toxin accumulation (Kim et al., 2015) and drought tolerance in vetch (De la Rosa et al., 2020). However, it is difficult to demonstrate the function of candidate genes due to the lack of a robust transformation and plant regeneration system (Ford et al., 2008;Nguyen et al., 2020). Preliminary research by Maddeppungeng (2006) showed that vetch cells are able to be transformed as evident by green fluorescent protein (GFP) expression in callus but no regeneration of plants from these transformed cells was reported. Maddeppungeng also reported the formation of embryogenic callus and subsequent plant regeneration from epicotyl explants (Maddeppungeng, 2006) but this has not been confirmed by others. Often regeneration of transformed cells is a bottle neck in many plant species, especially legumes (Somers et al., 2003). Although this can often be overcome by investigating different varieties, tissue types, media components and culture conditions, it is a very time consuming and laborious process which is by no means guaranteed of success.

An alternative to regenerating transgenic plants is to produce transformed hairy roots using the bacterium *A. rhizogenes*. Examples of success include soybean (Cho et al., 2000;Kereszt et al., 2007;Cai et al., 2015), *V. hirsuta* (Quandt et al., 1993), *V. faba* and *Medicago truncatula* (Vieweg et al., 2004). Previously, transgenic vetch hairy roots were described by Tepfer (1990), but there was no detailed description of the method. Hairy root induction requires the infection of pluripotent cells with *A. rhizogenes*, transfer and incorporation of the transfer DNA (T-DNA) into the plant genome and expression of *rol* genes encoded on the T-DNA. This often leads to the formation of roots with a “hairy” highly branched phenotype and the loss of plagiotropism (Sarkar et al., 2018;Tong et al., 2018). Including a second vector carrying a T-DNA containing genes of interest in the bacteria can lead to cotransformation into the plant nuclear genome (Tepfer, 1990). Co-transformation efficiency of up to 88% was reported in *V. hirsuta* (Quandt et al., 1993) but often ranges from 20 - 80% in other plants (Kereszt et al., 2007). Considering the possibility of using *A. rhizogenes* transformation, we aimed to develop a reliable, high-efficiency and time-saving protocol for hairy root transformation to facilitate functional gene analysis in vetch.

In this study, the hypocotyl region of five-day old seedlings were infected with *A. rhizogenes*. 28 days after inoculation of either *in vitro* or soil grown seedlings, about 24% or 43% respectively of infected seedlings produced transformed hairy roots. *In vitro,* the stable expression of a co-transformed reporter GUS was about 79%. In summary, transgenic hairy roots were readily obtained within a month, sufficient to facilitate functional gene analysis in vetch.

## MATERIALS AND METHODS

### Plasmid construction for GUS overexpression

The plasmid pMDC163 was modified by inserting a 35S CaMV promoter from pDONR207 by using a LR reaction (Invitrogen). The resulting plasmid: pMDC163::35S::GUS was transformed into *A. rhizogenes* strain K599 *via* electroporation. The presence of the plasmid in *A. rhizogenes* was confirmed by PCR detection by using oligonucleotide primers GUS_F and GUS_R that amplified a 552 bp product (**Table 1**).

**Table 1.**
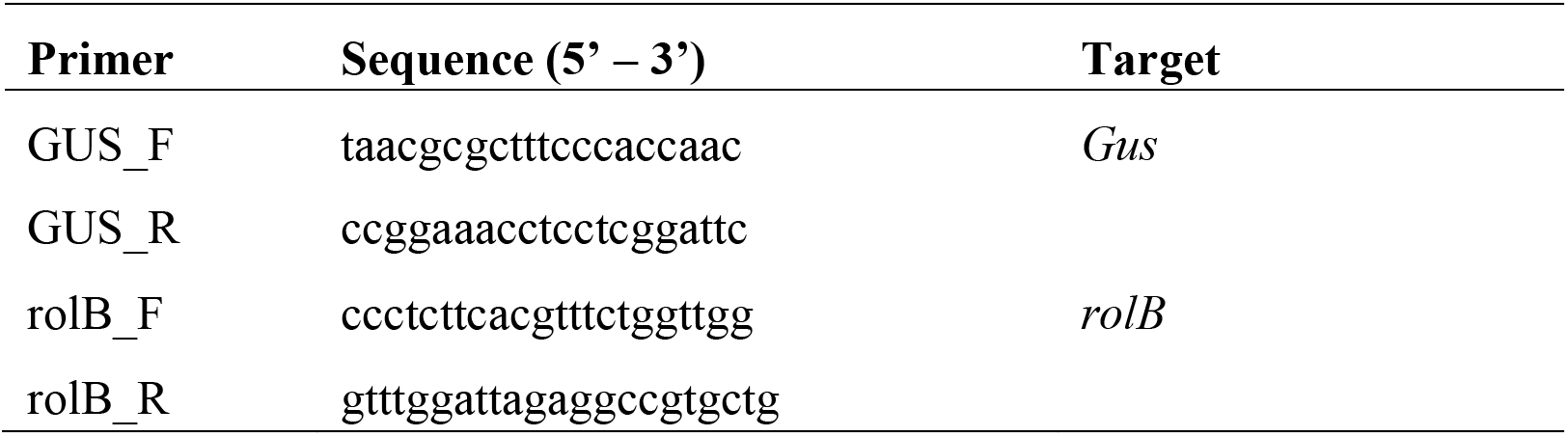
Oligonucleotide PCR primers used in this study.

### Preparing *A. rhizogenes* culture

Two cultures, *A. rhizogenes* K599 and K599 containing the plasmid pMDC163::35S::GUS were stored at −80°C. Prior to vetch transfection, the bacteria were streaked onto a LB plate or LB plate + 50 mg/L kanamycin to select for the presence of pMDC163::35S::GUS. A single colony was then inoculated into a 3 mL LB liquid culture and grown overnight at 30°C. The next day, 1 mL of the culture was mixed with 200 μl of 80% glycerol, and 200 μl of the mixture was spread onto a LB plate. The bacteria were cultured overnight in the dark at 30°C with the appropriate antibiotic in the media, if required.

### Vetch transformation on soil

Vetch seeds (variety Studenica) were sown 2 cm apart in a tray (30 x 50 x 15 cm) containing a mixture of sterile sand and cocopeat (1:1 in volume) and covered with a 1 cm layer of vermiculite. The seedlings were then grown under halogen lights (1000 lux) with 16 hr photoperiod, at 25°C (David et al., 2017). Five days after sowing, a small paste of *A. rhizogenes* was picked from the plate by using a 27G 1/2 (0.4.13 mm) needle and applied into the hypocotyl region of seedlings. The bacteria were pushed into the middle of the hypocotyls by inserting the needle 3 times. Next the infected regions were covered with vermiculite and watered with mist from a spray bottle. A transparent cover was placed over the tray to maintain high humidity. Water was sprayed regularly to keep the vermiculite layer moist. Nine days after infection (DAI), adventitious roots emerged from the wounded sites and were trimmed off to promote the development of hairy transformed roots. 23 DAI, hairy roots were observed and harvested for further analysis.

### Vetch transformation *in vitro*

Vetch seeds were surface sterilized with 70% EtOH for 1 min, followed by 5% NaOCl for 20 mins with vigorous agitation, and then rinsed with sterilized reverse osmosis water for 5 min and repeated 4 more times. The seeds were then sown on RGM_NoSuc medium (**Table 2**) and stratified at 4°C for 3 days before growing at 25°C in the dark for 2 days. Five days later, *A. rhizogenes* was introduced into the hypocotyl region similarly to the on-soil method described above. After infection, the infected seedlings were cultured on filter paper on a petri dish with RGM_NoSuc medium (**Table 2**) for about 23 days in the dark at 25°C. 23 DAI, the emergent hairy roots could be distinguished from wild type roots by their branching phenotype and were sub-cultured on RGM_3xSuc medium (**Table 2**) with added 25 mg/L meropenem (Sigma Aldrich). The hairy roots were sub-cultured every 4 weeks on the same medium and were cultured in the dark at 25°C.

**Table 2.** Medium formulation

| Component | Sigma Aldrich<br>Cat. Number | RGM_NoSuc | RGM_3xSuc |
| --- | --- | --- | --- |
| MS (Murashige and Skoog) Basal | M5519 | 2.2 g/L | 2.2 g/L |
| 1000x MS vitamins | M3900 | 1 mL/L | 1 mL/L |
| myo-inositol | I7508 | 200 mg/L | 200 mg/L |
| MES hydrate | M2933 | 0.5 g/L | 0.5 g/L |
| Sucrose |  | 0 g/L | 30 g/L |
| Gelzan | G1910 | 3 g/L | 3 g/L |
| Adjust pH with 1M KOH |  | 5.7 | 5.7 |

### PCR confirmation of transgenic hairy roots

The hairy root phenotype is caused by expression of *rol* genes from the T-DNA of pRi2659 after incorporation into the vetch nuclear genome. rolB_F- and rolB_R-specific primers which amplified a 381 bp PCR product were used in PCR amplification to detect the *rolB* gene in transgenic hairy roots (**Table 1**).

### Histochemical β-glucuronidase (GUS) expression assay for transgenic hairy roots

Hairy roots were incubated with substrate solution, 2 mM XGluc (cat. G1281C1, Gold Biotechnology), 0.5 mM K_3_Fe_6_, 0.5 mM K_4_Fe_6_, 50 mM NaPO_4_ pH 7.2 and 0.1% (v/v) Triton X-100 at 37°C overnight. The roots were then washed with 100% EtOH 3 times and incubated in EtOH for at least 1 hr prior to observation under a stereo microscope (Olympus SZ2_ILST). Hairy roots that were PCR negative for the GUS vector were used as a negative control for staining.

## RESULTS

### Hairy root induction on soil

Five-day old soil grown seedlings were infected with *A. rhizogenes* by pushing the bacterial inoculum into the hypocotyl region using a small gauge needle. High humidity was maintained at the wound site by keeping a vermiculite layer moist all times (**Figure 1A**). 9 days after inoculation (DAI), callus and adventitious roots were observed at the wound site (**Figure 1A, E**) but they did not have the required hairy root phenotype. In the negative control plant, wounds were introduced but without the bacteria and there was no callus formation (**Figure 1D**). However, a few plants did show adventitious roots protruding from the wounded sites (not shown). Taken together, the adventitious roots 9 DAI might be just the result of wounding, so these roots were trimmed to promote the growth of transgenic hairy roots. 23 DAI, roots protruding from the callus at the infected sites showed a typical hairy root phenotype: highly branching and showing the loss of plagiotropism (**Figure 1A, G**). These roots were absent in the negative control plant (**Figure 1F**). PCR detection was performed to confirm the presence of the *rol* genes in the hairy roots. The PCR results showed the presence of *rolB* gene in the hairy roots tested but there were no PCR products in the non-hairy roots that came from the same plant (**Figure 1H**). In this experiment, 43% of seedlings were confirmed to be transgenic hairy roots (**Table 3**).

**Table 3.**
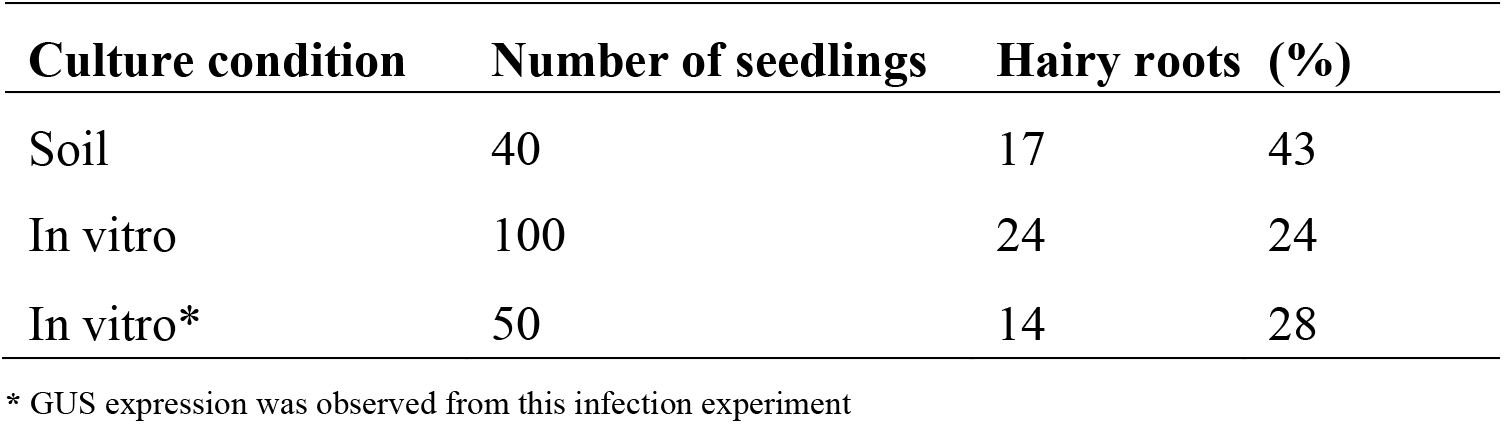
Percent of seedlings with hairy roots after transfection with *A. rhizogenes* K599.

**Figure 1.**
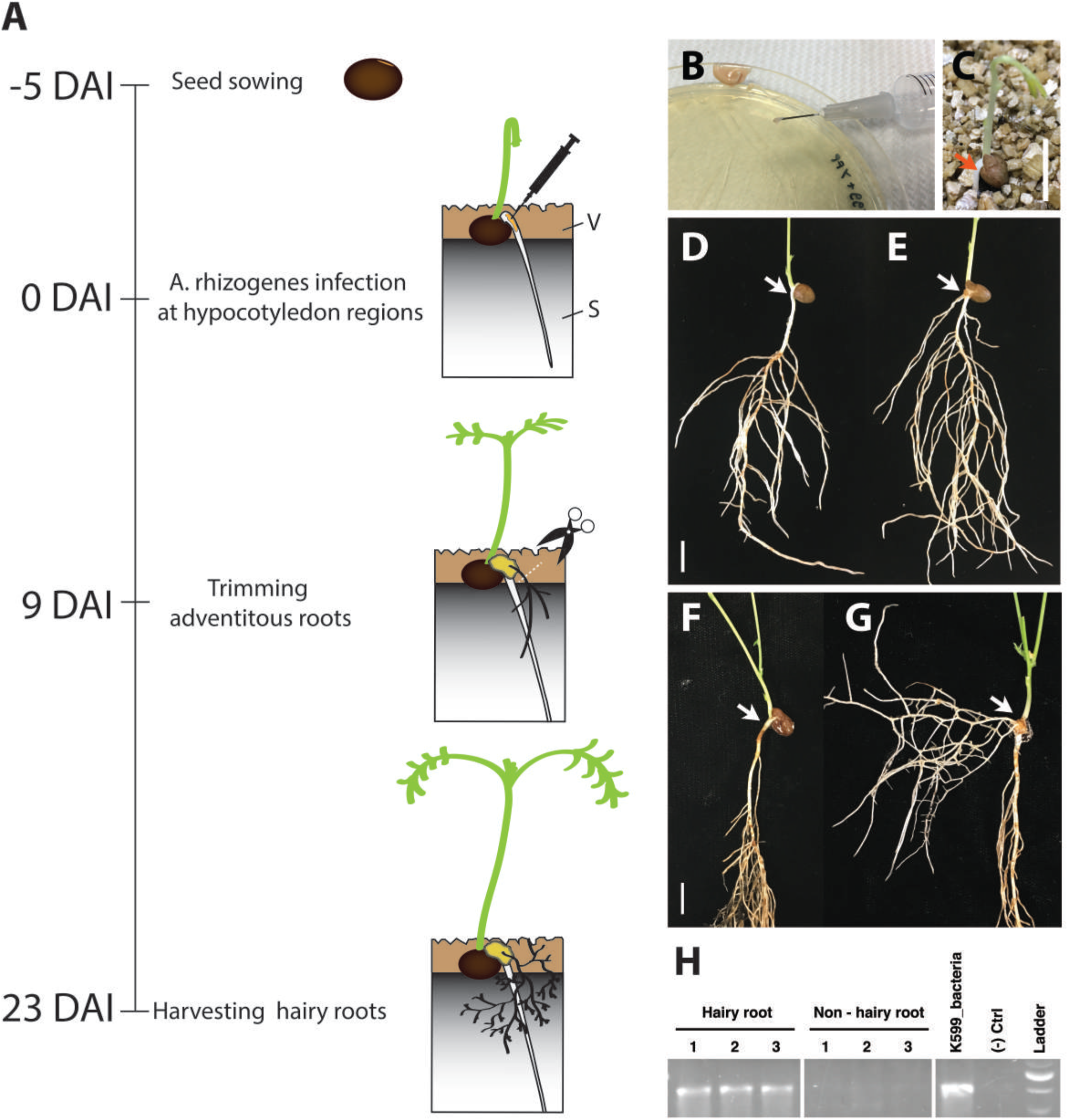
Hairy root induction in vetch grown on compost by *A. rhizogenes* strain K599. (A) Timeline for hairy root induction on soil by *A. rhizogenes*. On day 1, vetch seeds were sown on top of a mixture of sand and cocopeat (1:1), and 1 cm of vermiculite was placed on top to cover the seeds. Seeds were allowed to germinate for 5 days. On day 5, seedling hypocotyl regions were infected with *A. rhizogenes*. 9 days after infection (DAI), adventitious roots were observed at the wounded region and were trimmed off to promote the growth of hairy roots. 23 DAI, hairy roots were observed. (B) *A. rhizogenes* were grown on a LB plate and scraped into a paste which was pushed into the hypocotyl region of plants by stabbing a 27G ½ needle. (C) 5-day old seedling with the red arrow showing the position to be inoculated using the stabbing method. (D) A 9 DAI control seedling that was stabbed with a needle containing no bacteria (negative control). No adventitious root or callus appeared at the wounded site. (E) A 9 DAI seedling that was inoculated with *A. rhizogenes* showed the formation of callus and adventitious roots at the wounded site and the adventitious roots had a wild type phenotype; adventitious roots from (E) were later trimmed as they were not transgenic. (F) A negative control 23 DAI plant showed no callus or adventitious roots at the wounded sites. (G) At 23 DAI the inoculated plants showed the formation of callus and hairy roots which were highly branched and had lost plagiotropism. (H) The *rolB* gene was detected by PCR amplification in hairy roots but not in the nonhairy roots from the same plants (3 biological replicates are shown). DAI = day after the infection, V = vermiculite, S = sand and cocopeat (1:1, volume:volume). Scale bar = 1 cm.

### Hairy root induction *in vitro*

Sterilized seeds were sown on RGM_NoSuc media and stratified for 3 days before germination in the dark at 25°C. Then *A. rhizogenes* was infected into the hypocotyl region similarly to the on-soil method, but in a sterile laminar flow hood (**Figure 2A**). To avoid the infected sites contacting the media, the infected sites were placed onto filter paper while the roots were still touching the media (**Figure 2A, B**). The seedlings were then cultured in the dark on the same media. Callus formation was observed at the wounded sites (**Figure 2C**) and 23 DAI hairy roots were observed (**Figure 2D**). Compared with wild type roots, hairy roots showed very early lateral root formation and vigorous growth. As a result, hairy roots were much more branched when compared with wild type (**Figure 2E**). Moreover, due to the loss of geotropism, hairy roots not only penetrated into the media but also grew upwards (not shown). In this experiment, the induction of hairy roots occurred in about 24% of the infected seedlings (**Table 3**). Similar to hairy root induction on soil, 1 - 3 hairy roots were observed from each inoculated seedling. Each of these hairy roots were considered to result from an independent transformant.

**Figure 2.**
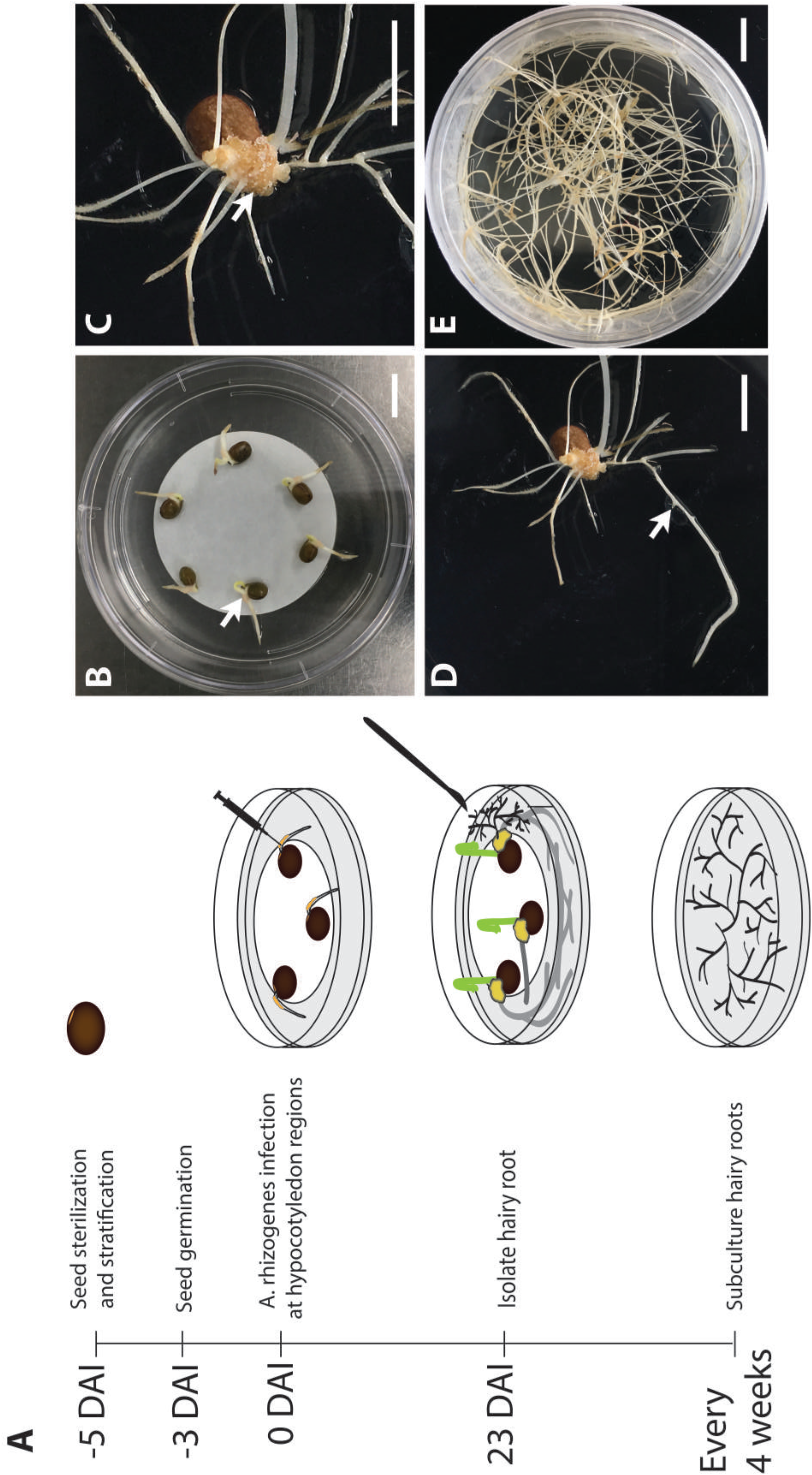
Hairy root induction in vetch by *A. rhizogenes* K599 *in vitro*. (A) Timeline for *in vitro* hairy root induction by *A. rhizogenes*. On day 1, sterilized seeds were sown on a plate containing RGM_NoSuc and stratified at 4°C in the dark. On day 3, seeds were placed at 25°C in the dark for germination. On day 5, the seedling hypocotyl region was inoculated with *A. rhizogenes* by using the stabbing method and then cultured on RGM_3xSuc medium with a filter paper to prevent bacterial overgrowth. 23 days after the infection (DAI), hairy roots were isolated and sub-cultured onto RGM_3xSuc medium with 25 mg/L meropenem. Hairy roots were sub-cultured every 4 weeks onto fresh medium. (B) Inoculation of seedling hypocotyl regions (white arrow) with *A. rhizogenes*. (C) 23 DAI callus developed from the inoculated region (white arrow) and (D) a hairy root was observed on the surface of the callus with its lateral roots forming at a very early stage (white arrow), which could help to distinguish hairy roots from wild type roots. (E) Hairy roots 45 days after the inoculation were highly branched. DAI = day after the infection. Scale bar = 1 cm.

### Chimeric expression pattern of the GUS transgene

To determine the co-transformation efficiencies of the T-DNAs from pRi2659 and pMDC163::35S::GUS by *A. rhizogenes,* seedlings were inoculated *in vitro* and hairy roots were used in GUS assays. In this experiment, 14 hairy roots were obtained from 50 seedlings (**Table 3**). Of 14 roots, 86% showed GUS expression (**Table 4**). However, different patterns of the GUS expression in the lateral hairy roots were observed (**Table 4**). Out of the 12 hairy root lines analyzed, 6 contained GUS activity in both the primary and lateral roots (**Figure 3D**), 5 showed GUS activity in the lateral roots (**Figure 3F**), and 1 showed GUS activity in the primary root and not in the lateral roots (**Figure 3E**).

**Table 4.** GUS expression patterns in transformed hairy roots.

| GUS localization | Number of hairy root(s) | (%) |
| --- | --- | --- |
| Primary and lateral root | 6 | 43 |
| Primary root only | 1 | 7 |
| Lateral root only | 5 | 36 |
| Not detected | 2 | 14 |
| Total | 14 | 100 |

**Figure 3.**
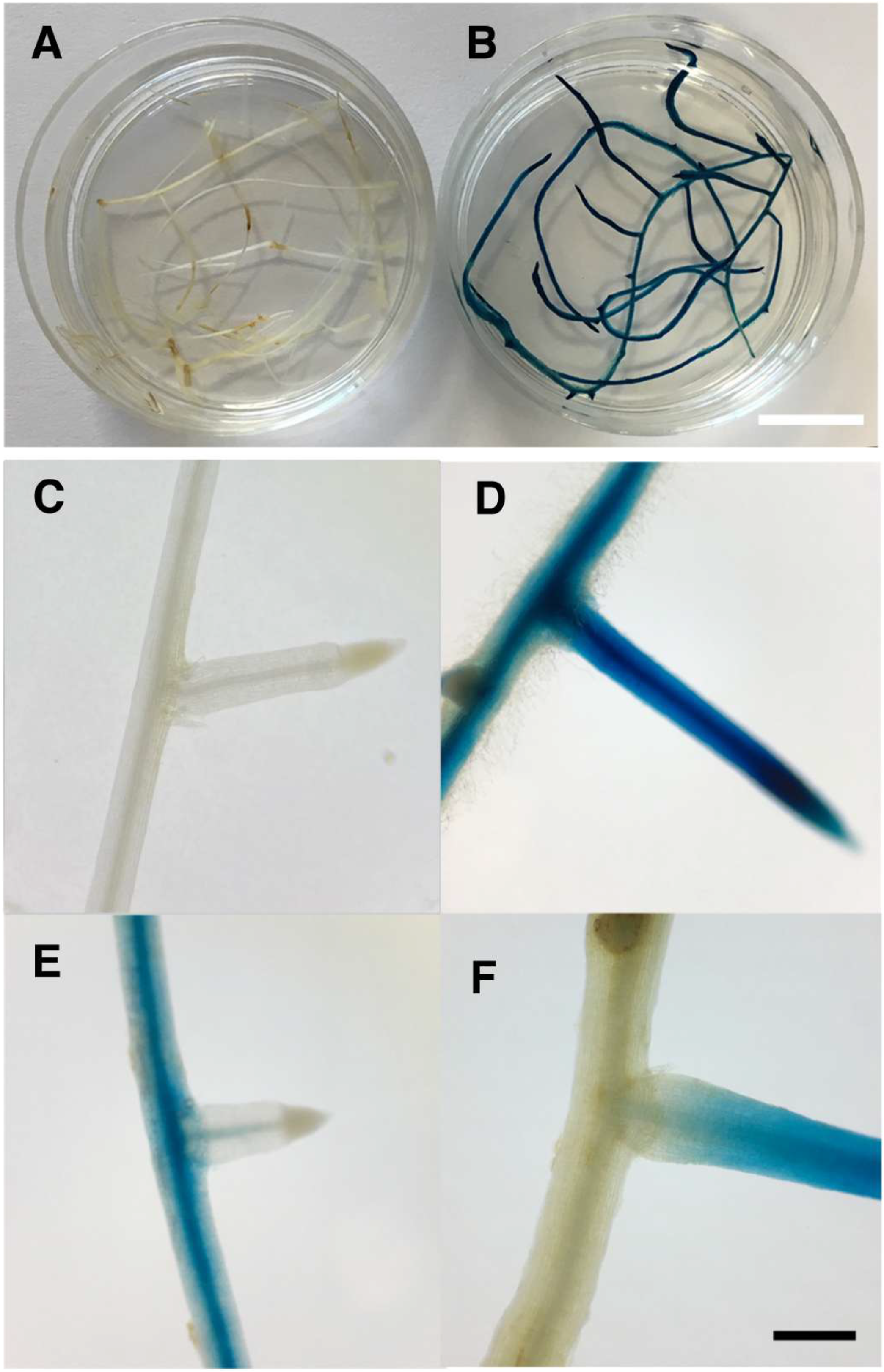
Transformed hairy roots with GUS activity. (A) Hairy roots from seedlings infected with *A. rhizogenes* K599. (B) Hairy roots from a seedling infected with *A. rhizogenes* that contained a plasmid containing the GUS reporter gene. (C) A hairy root without GUS activity was used as a control. (D, E, F) showed different patterns of GUS expression in hairy roots depending on the developmental stage at which the root stem cells were transformed. (D) GUS expression both in the primary and lateral hairy roots. (G) Loss of GUS activity in lateral hairy root. (H) GUS expression in the lateral root. (A, B) scale bar = 1 cm; (E, F, G, H) scale bar = 3 mm.

### The deterioration of vetch hairy root growth under prolonged culture *in vitro*

Hairy roots were maintained by sub-culturing onto a new RGM_3xSuc medium every 4 weeks. For each subculture, a 5 cm segment of hairy root which contained about 5 – 10 lateral roots were transferred onto new RGM_3xSuc medium. After a 4-week-growth period, we observed extensive root growth that covered the surface of the media in the 9 cm petri dish as shown in **Figure 4A**. Vigorous hairy root growth was maintained for about 4 months. After that, a significant reduction in root growth was observed: the roots turned brown and new established lateral roots were thin and slow growing with a swollen root tip (**Figure 4B, C)**.

**Figure 4.**
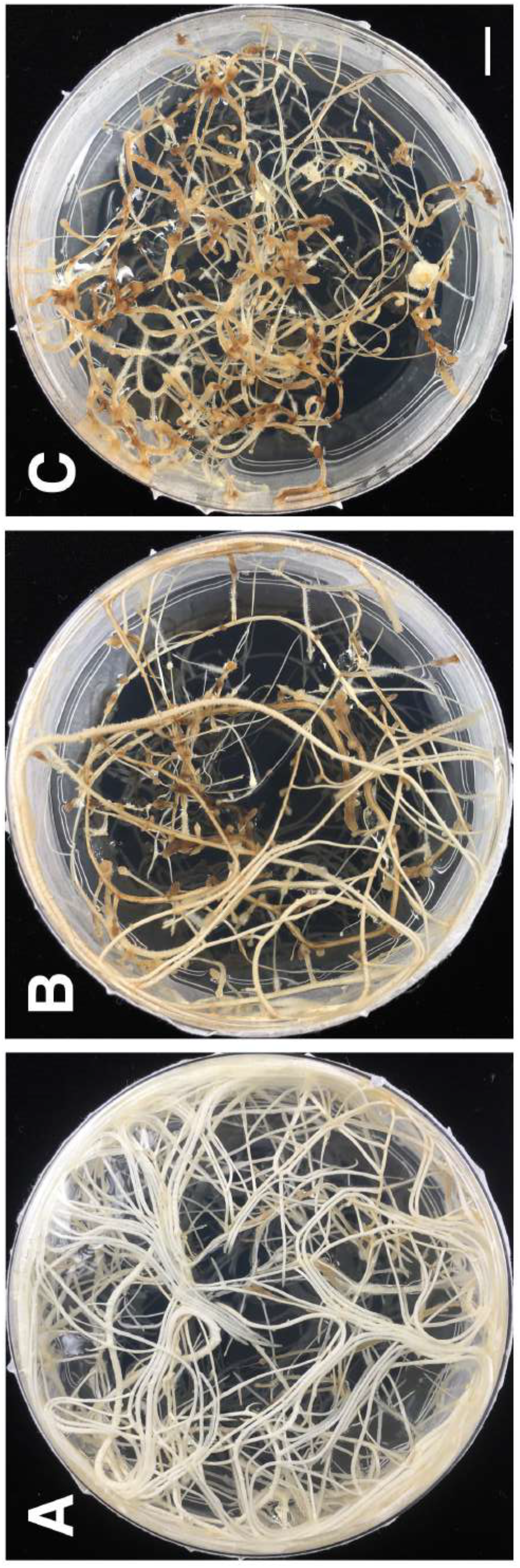
Deterioration of hairy root growth under prolonged culture *in vitro*. (A) 2-month-old hairy roots. (B) 5-month-old hairy roots. (C) 6-month-old hairy roots. Scale bar = 1 cm.

## DISCUSSION

### The difference in transformation efficiency between soil and *in vitro* grown seedlings

The transformation efficiency of soil grown seedlings was 43% but this was reduced to 24% in the *in vitro* grown seedlings (**Table 3**). This reduction was possibly due to the developmental age of the seedlings when transfected or due to the culture conditions. At the time of transfection, seedlings on soil were 5 days old after sowing and large in size, while the *in vitro* grown seedlings were smaller and the roots were about 1-2 cm in length (**Figure 2B**). Also, the infection sites of soil grown seedlings did not show any necrosis symptoms when compared to the *in vitro*-grown seedlings possibly as the drainage and ventilation were better (data not shown). We observed extensive bacterial growth on the media after infection when the infected tissue was in contact with the medium in early experiments (data not shown). In these early experiments, the bacteria would over-grow and the tissues became wilted leading to cell death. No callus formation or hairy roots were observed on these bacterial overgrown tissues. To minimize this bacterial overgrowth in subsequent *in vitro* infections, the infected seedlings were laid on sterile filter paper and cultured on media without sucrose. It was also important to position the seedlings so that the infected sites were on the filter paper, while maintaining the existing roots in contact with the media (**Figure 2A, B**). In this way we could effectively stop the bacterium over-growing and eliminate the need to treat seedlings with antibiotics as required in other protocols (Böttinger et al., 2001;Olhoft et al., 2007). With the *in vitro* method, due to the lack of sucrose and no light during the 28-day growth period, the plants grew slowly, and the roots were thinner when compared to soil grown seedlings. The developmental stage of the seedlings and the stressful *in vitro* culture conditions, which were necessary to control the bacteria growth but did not favor the plant growth, together may have compromised the transformation efficiency. However, with this *in vitro* method, infections could be quickly performed 5 days after germination and no extra steps were required until the hairy roots developed.

In both the soil and *in vitro* transformation systems, the trimming of adventitious roots at an early stage facilitated hairy root growth. While not reported here, removal of the main roots can be performed and the hairy roots support the shoot growth (Kereszt et al., 2007). The effect of removing the primary root to promote hairy root development is worthwhile testing in the future. Presumably trimming of the main root could overcome the dominant effect of the primary root and hence promote the growth of lateral roots and hairy roots from the callus.

### Chimeric GUS expression in transgenic hairy roots

Adventitious and lateral root development often starts with a single cell that acquires pluripotency, and it is often a pericycle cell that is adjacent to the xylem pole. This pluripotent cell further develops into a multi-cellular stem cell niche that includes a quiescent center that gives rise to different cell layers of a root (Shemer et al., 2015;Radhakrishnan et al., 2018). During the period from pluripotent acquisition to the formation of the quiescent center, depending at what developmental stage is transformed by the *Agrobacterium,* may explain the holistic or chimeric transgenic expression pattern in the resulting root. Only the transformed cell and its daughter cells will contain the transgene, therefore, to obtain the holistic transgenic expression, the transformation event must happen before or just after the pluripotent acquisition of a single cell. As seen in our experiment, the GUS expression pattern observed in both the primary and lateral hairy roots may be explained by the original cell that was infected. For example, if a pluripotent cell divides to give rise to a stem cell niche and one of these daughter cells was transformed, then only the cells derived from this daughter cell would show GUS activity. **Figure 3E** shows an example of a daughter cell that was transformed and gave rise to transformed vascular cells but these vascular cells (xylem and phloem) could not differentiate into lateral roots, such that GUS activity was not inherited to the lateral roots. In contrast, other studies revealed lateral root formation starting with pluripotency acquisition of a pericycle cell adjacent to xylem pole (Atta et al., 2009;Shemer et al., 2015). Hence, if this pericycle cell layer had been transformed with GUS, then the lateral roots would inherit this activity as seen in **Figure 3F**. As the pericycle cell layer was thin, GUS activity was low in the primary root but appeared strongly in lateral roots as all the cell layers were now expressing GUS (**Figure 3F**).

Our transformation method efficiently produces transgenic hairy roots by infecting vetch hypocotyls with *A. rhizogenes*. Our result suggests that the targets for transformation are the pericycle cell and the derived stem cell niche. However, it has been shown that other tissue types can be transformation by *A. rhizogenes,* for example, hairy roots were obtained from infected epicotyls of *V. hirsuta* (Quandt et al., 1993), or leaf disks of *Duboisia leichhardtii* (Mano et al., 1989) where there are no pericycle cells present. Whether chimeric transgene expression occurs in transgenic roots derived from these other tissue types is unclear. Currently it is unknown if other tissue types of vetch can be infected by *A. rhizogenes* to produce hairy roots.

In brief, when using our transformation method which targets the cells in the hypocotyl region, to reduce chimeric expression of transgenes in the derived hairy roots, it is important to perform the transfection as early as possible when the stem cell niche has not formed in the hypocotyl. This study also emphasizes the need to confirm transgene expression in the hairy roots before using the roots in subsequent experiments as we show that not all the hairy roots inherit transgene expression.

### Deterioration of hairy root growth under *in vitro* conditions

It was not surprising to observe in vetch that hairy root growth decreased after a few months of *in vitro* culture as vetch is an annual plant. The underlying molecular causes of the decreased growth are unclear but may be due to epigenetic changes that are inherited over prolonger culture periods. This may suggest that the hairy root material from vetch and other annual plants should be used within 3 months after infection to ensure the highest quality root material.

### Conclusions

This efficient and time-saving hairy root induction method in vetch using *A. rhizogenes* strain K599 bacteria will greatly facilitate gene expression and functional studies. However, due to the nature of chimeric transgenic root formation, lateral roots should be confirmed to contain and express the transgene.

## Author contributions

IRS conceived the project. VN conducted the experiments, interpreted the results and wrote the paper. IRS interpreted the results and edited the paper.

## Acknowledgments

The authors would like to thank the Hermon Slade Foundation (Grant number HSF1707) and The University of Adelaide for funding awarded to VN and IRS. The authors would like to thank Jeremy Timmis for assistance with editing the manuscript.

## Notes

### Competing Interest Statement

The authors have declared no competing interest.

